# Chemokine receptor expression defines a trajectory from monocytes to mature macrophages

**DOI:** 10.1101/2025.10.27.684776

**Authors:** Heather Mathie, Laura Medina-Ruiz, Fabian Schuette, Heba Halawa, Zuzanna Pocalun, Elise Pitmon, John Cole, Marieke Pingen, Gerard J Graham

## Abstract

CCR1, CCR2 and CCR5 direct recruitment of monocytes and macrophages in inflammation. However, the discrete role for each receptor in monocyte/macrophage biology remains poorly understood, with previous reports citing receptor redundancy. Using transcriptomic approaches to examine inflammatory chemokine receptor expression on lung interstitial macrophage populations, we demonstrate that interstitial macrophages can be divided into three distinct subsets, each of which express specific patterns of chemokine receptors, and that there are dynamic changes in chemokine receptor expression as macrophages differentiate from monocytes in the lung. Furthermore, macrophages expressing different combinations of chemokine receptors are transcriptionally distinct, suggesting non-redundant functions for CCR1, 2 and 5. Finally, we examined changes in macrophage chemokine receptor expression *in vitro* after treatment with varied TLR ligands, and show that CCR1 is specifically increased in response to bacterial but not viral ligands. Our data provide compelling evidence that macrophage chemokine receptor expression is not redundant, but specific and malleable in response to discrete inflammatory stimuli.

## INTRODUCTION

Monocyte-derived macrophages are an essential component of the inflammatory response and are implicated in pathogen, and cellular debris, removal and wound healing(Wynn et al., 2013). The recruitment of monocytes to inflamed sites, and the interstitial movement of their macrophage progeny within the inflamed tissue, are regulated mainly by members of the chemokine family(Griffith et al., 2014). Chemokines are characterised by a conserved cysteine motif and divided into CC, CXC, XC and CX3C subfamilies accordingly(Rot and von Andrian, 2004; Zlotnik and Yoshie, 2000). They mediate their effects through cognate 7-transmembrane spanning receptors which are named according to the subfamily of chemokine to which they bind i.e. CCRs, CXCRs, XCR and CX3CR(Bachelerie et al., 2014). Broadly speaking, chemokines, and their receptors, can be defined as being inflammatory or homeostatic according to the contexts in which they function(Zlotnik and Yoshie, 2000). Monocyte and macrophage dynamics during the inflammatory response are predominantly controlled by inflammatory CC-chemokines and their cognate receptors.

We are interested in defining chemokine receptor involvement in monocyte and macrophage dynamics at inflamed sites. Literature in this area is confusing with reports suggesting significant redundancy of chemokine receptor expression patterns on both monocytes and macrophages(Mantovani, 1999; Mantovani, 2018; Proudfoot, 2002). Our focus has been on a single chromosomal locus incorporating the receptors CCR1, CCR2, CCR3 and CCR5(Dyer et al., 2019; Medina-Ruiz et al., 2022). Of these receptors, CCR1, CCR2 and CCR5 (henceforth inflammatory chemokine receptors or iCCRs) are clearly implicated in monocyte and macrophage function at inflamed sites. Using novel mouse models, we have demonstrated that, in 98% of inflammatory monocytes, CCR2 is the only iCCR that is significantly expressed(Dyer et al., 2019; Medina-Ruiz et al., 2024; Medina-Ruiz et al., 2022). Mice bearing a compound deletion of the iCCR locus show the anticipated profound monocytopenia previously reported in CCR2 deficient mice confirming an essential, and nonredundant, role for CCR2 in monocyte egress in the bone marrow(Dyer et al., 2019). Our data also demonstrate that CCR2 is central to recruitment of monocytes from the circulation to the tissue. Thus, the issue of redundancy of iCCR expression is not apparent in the context of monocytes, which are fully reliant on CCR2. In contrast to CCR2, expression of CCR1 and CCR5 becomes apparent once monocytes start to differentiate towards macrophages within peripheral tissue(Medina-Ruiz et al., 2022) and here things become much more complicated. To study iCCR biology, we previously generated compound transgenic reporter (REP) mice, with a recombineered bacterial artificial chromosome (BAC) inserted into the genome, in which the coding sequence of each of the iCCR genes was replaced with sequences encoding spectrally distinct fluorescent proteins. Therefore, in REP mice, as an iCCR gene is expressed, expression of a corresponding fluorescent protein will also occur. Using these REP mice, we have shown that individual macrophages, within both resting and inflamed tissues, can express all combinations of CCR1, CCR2 and CCR5(Medina-Ruiz et al., 2022).

The purpose of the present study was to examine the dynamics of iCCR expression by macrophages, how dynamics alter as macrophages differentiate from monocytes, and to determine whether macrophage subsets display chaotic or specific patterns of receptor expression. In addition, we aimed to determine if iCCRs have a redundant or non-redundant role in the macrophage response to varied inflammatory stimuli. Here we have used transcriptomic approaches to demonstrate that resting lung interstitial (IM) macrophages expressing different combinations of iCCRs are transcriptionally distinct. Furthermore, we demonstrate, using single cell RNA sequencing, that monocyte to macrophage differentiation is characterised by transition through an early monocyte-derived IM population, the largest proportion of which express CCR1, CCR2 and CCR5. We propose that this early recruited macrophage population, which we have termed Retnla+ macrophages, express a combination of iCCRs to allow them to respond to a range of environmental cues. As macrophages terminally differentiate, they tailor iCCR expression along divergent trajectories, depending on the phenotype of terminally differentiated cell, thereby altering iCCR expression to align with macrophage function. Our data shed new light on the nature of monocyte-to-macrophage differentiation and demonstrate that iCCR expression on macrophages is not redundant or chaotic but specific and malleable in response to discrete inflammatory agents.

## RESULTS

### Bulk RNA sequencing demonstrates that IMs expressing different combinations of iCCRs are transcriptionally distinct

Bulk RNA-sequencing was performed on cells sorted from MHCII hi and MHCII lo IM populations based on iCCR expression patterns, in order to determine if cells expressing different combinations of iCCRs are transcriptionally distinct and thus possess discrete functionality. For this analysis, different iCCR-expressing macrophage populations were sorted from REP mice. The gating strategies are presented in Supplementary Figure 1A and Figure 1A. Macrophages were gated first on MHCII expression, to delineate MHCII hi versus MHCII lo macrophages, then gated on iCCR expression patterns. iCCR populations sorted from the MHCII hi gate included: CCR5+ cells, CCR1+CCR5+ cells, CCR2+CCR5+ cells and CCR1+CCR2+CCR5+ cells (Figure 1Ai). MHCII lo IMS were also sorted based on iCCR expression patterns. Cells sorted from the MHCII lo gate included: iCCR-ve cells, CCR2+ cells, CCR5+ cells, and CCR1+CCR5+ cells (Figure 1Aii). Cells from each of these gates were collected for RNA sequencing. PCA analysis of the MHCII hi IMs revealed that each population clustered separately on a PC-1 vs PC-2 scatterplot, demonstrating that there are transcriptional differences between each of the iCCR expressing populations (Figure 1Bi). This is further demonstrated by the heatmap in Figure 1Bii, which shows significantly differentially expressed genes between MHCII hi IMs expressing different iCCR combinations. This analysis suggests that the CCR1+CCR2+CCR5+ population has the most highly distinct gene expression profile of all MHCII hi IMs, but that all populations are unique (Figure 1Bii).

**Figure 1:**
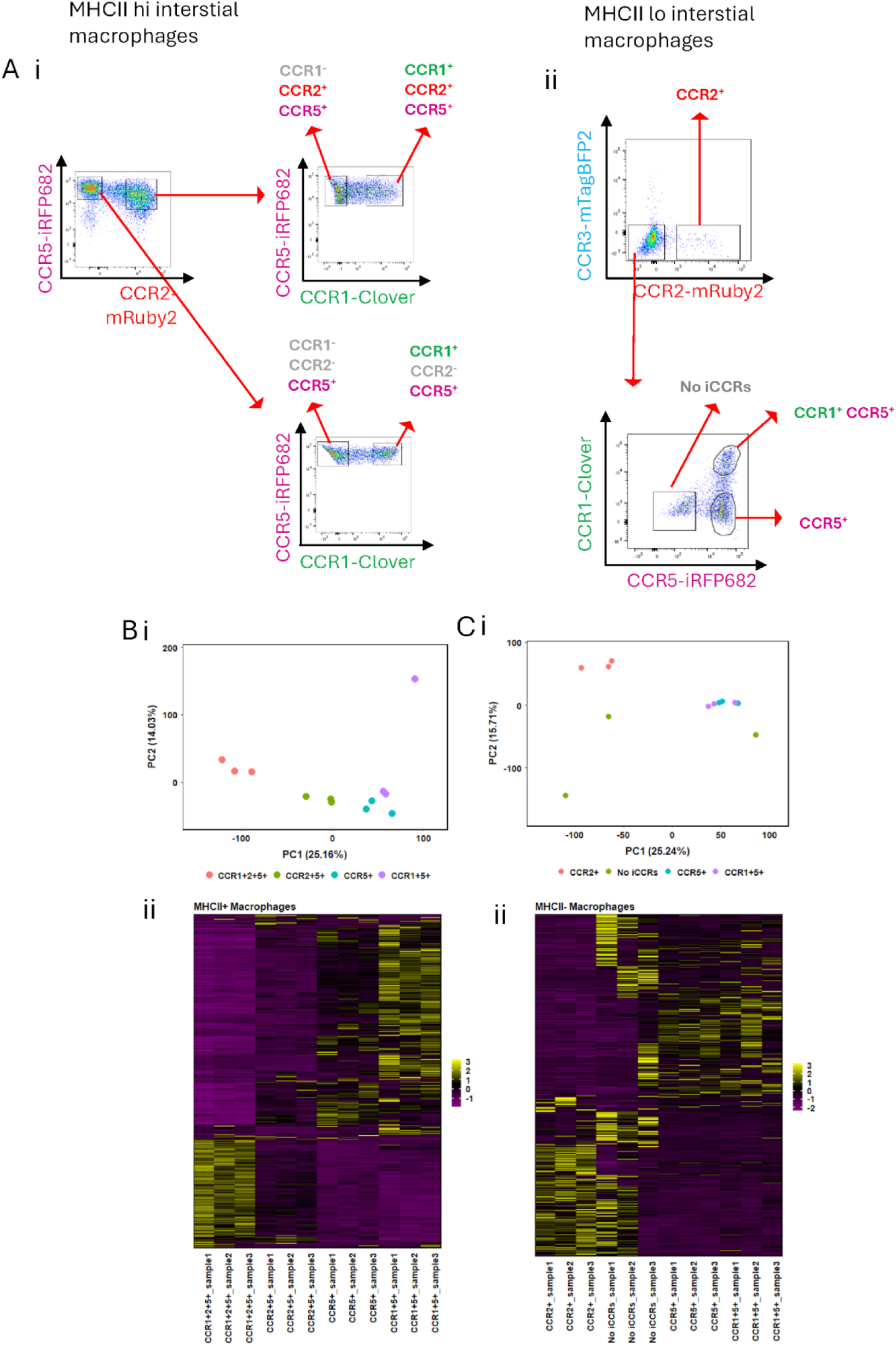
Bulk RNA-Sequencing demonstrates that lung IM populations expressing different iCCR combinations are transcriptionally distinct. (A) (i) Flow cytometry plot showing the iCCR expression patterns of MHCII hi macrophages. (ii) Flow cytometry plot showing the iCCR expression patterns of MHCII lo macrophages. (B) (i) Principal component analysis scatter plot showing PC1 versus PC2 for MHCII hi macrophages expressing various combinations of iCCRs. Percentage variance is displayed in brackets for each principal component. (ii) Heatmap showing significantly differentially expressed genes between MHCII hi macrophages displaying differing iCCR expression patterns. (C)(i) Principal component analysis scatter plot showing PC1 versus PC2 for MHCII lo macrophages expressing various combinations of iCCRs. Percentage variance is displayed in brackets for each principal component. (ii) Heatmap showing significantly differentially expressed genes between MHCII lo macrophages displaying differing iCCR expression patterns. For heatmaps in B ii and C ii, significantly differentially expressed genes are displayed on the y-axis and were hierarchically clustered using Spearman distances with UPMGA agglomeration and mean reordering. Expression levels were row-scaled into z-scores. Colour represents expression level, with purple representing low expression and yellow representing high expression.

Analysis of MHCII lo IMs revealed that CCR2 expression was depleted compared to MHCII hi IMs (Figure 1Aii). In addition, MHCII lo IMs also contained a population negative for iCCR expression (Figure 1Aii). PCA analysis of MHCII lo IMs revealed that, while iCCR-ve cells and CCR2+ cells formed distinct clusters on a PC-1 vs PC-2 scatterplot, CCR1+CC5+ and CCR5+ cells clustered together (Figure 1Ci). A heatmap depicting significantly differentially expressed genes between MHCII lo IM populations confirms this observation (Figure 1Cii), demonstrating CCR2+ and iCCR-ve cells to have unique gene expression patterns, whereas the gene expression patterns for CCR1+ and CCR1+CCR5+ MHCII lo IMS was highly similar.

Essentially, the bulk RNA-Seq data shows that MHCII hi and MHCII lo IMs expressing various combinations of iCCRs are transcriptionally distinct (with the exception of CCR1+CCR5+ and CCR5+ MHCII lo IMs), suggesting discreet roles for iCCRs in the inflammatory response.

### Single cell RNA sequencing confirms that differential iCCR expression patterns mark distinct IM subsets

Next, single cell RNA sequencing was employed to investigate the transcriptional relationship between different iCCR-expressing IM populations in the resting lung. IMs were enriched from the lungs of REP mice(Medina-Ruiz et al., 2022) by sorting CD11b+ SiglecF− Ly6G− cells, in order to deplete alveolar macrophages, eosinophils and neutrophils from the analysis. The enriched population was then FACS-sorted based on the expression of CCR1 and CCR5 reporters (gating strategy for sorts is demonstrated in Supplementary Figure 1B). The resulting four populations of enriched cells (CCR1− CCR5+, CCR1+ CCR5+, CCR1+ CCR5−, CCR1− CCR5−) were separately hash-tagged using different TotalSeq-A antibodies in order to accurately identify the iCCR expressing populations during sequencing analysis. This step was important as transcript and protein expression levels do not always align for CCR1 and CCR5 making transcript levels unreliable as an indicator of receptor protein expression(Hariharan et al., 1999; Kaufmann et al., 2001). Hash-tagged samples were integrated together with an enriched, non-hash-tagged sample of CD11b+ SiglecF− Ly6G− lung cells.

Bioinformatic analysis revealed 16 clusters of cells, including NK-cells, dendritic cells, monocytes, macrophages, B-cells, granulocytes and stromal cells (Figure 2A). A full list of genes differentially expressed between clusters can be found in Supplementary Table 1. The data were then sub-setted to depict only monocytes and macrophages. Monocytes clearly separated into 2 distinct populations which could be delineated by Ly6C expression; the largest population consisting of Ly6C hi classical monocytes, with a smaller population depicting Ly6C lo non-classical monocytes. In the current study no trajectory could be identified between non-classical monocytes and IMs (data not shown) in the resting lung, therefore non-classical monocytes were removed from downstream analysis and the data re-scaled (Figure 2B). A full list of genes differentially expressed between the monocyte and macrophage clusters depicted in Figure 2B clusters can be found in Supplementary Table 2.

**Figure 2:**
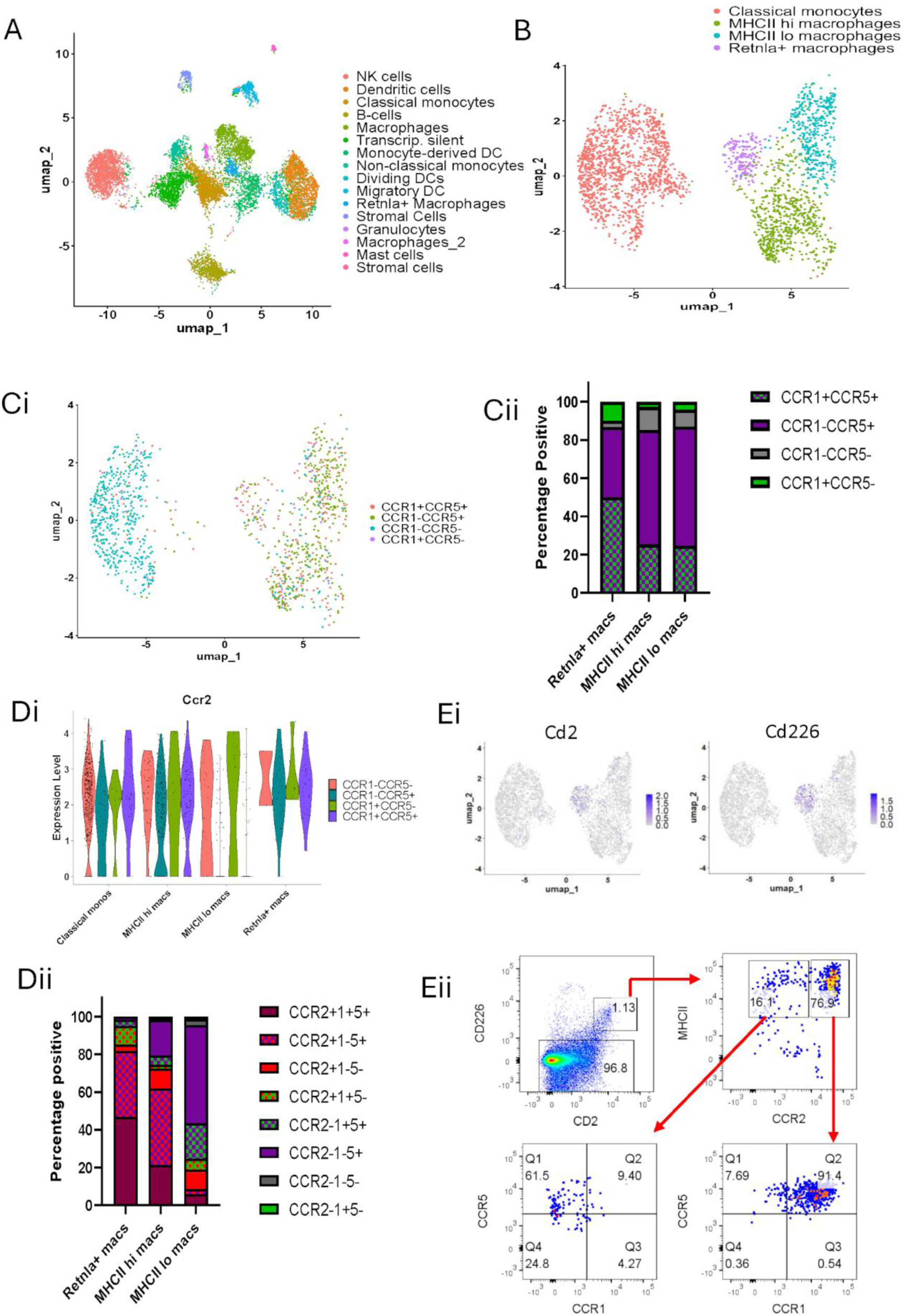
Single cell RNA-sequencing reveals that subsets of lung IMs have unique iCCR expression patterns and that iCCR expression is not random. (A) Uniform Manifold Approximation and Projection (UMAP) visualization of CD11b+ Ly6G− SiglecF− lung cells. (B) UMAP plot of data sub-setted from A depicting cells positive for monocyte or macrophage markers. Non-classical monocytes were also removed from the data-set. Colours represent unique clusters termed “classical monocytes”, “*Retnla+* macrophages”, “MHCII hi macrophages” and “MHCII lo macrophages”. (C)(i) UMAP displaying hash-tagged populations and how they correspond to the monocyte and macrophage clusters presented in B. Blue=CCR1-CCR5-, Red=CCR1+CCR5+, Green=CCR1-CCR5+, Purple=CCR1+CCR5-. UMAP plots in B and C have the same co-ordinates. (ii) Bar graph presenting the proportion of hash-tagged cells corresponding to each macrophage population. (D)(i) Violin plot showing the relative expression of *Ccr2* by hash-tagged populations across the four clusters presented in B. (ii) Bar graph presenting combinatorial iCCR expression across the three macrophage populations presented in B. Cells were considered positive for *Ccr2* if relative expression was >1. Feature plots depicting the expression of *Cd2* and *Cd226* within the single cell RNA-seq dataset. UMAP co-ordinates are identical to the plot shown in Figure 2 B. (i) Antibodies against CD2 and CD226 were used to identify *Retnla+* macrophages by flow cytometry and assess iCCR expression in the CD2+ CD226+ population.

Similar to previous studies(Gibbings et al., 2017), we found that lung IMs can be separated into 3 distinct clusters based on gene expression patterns (Figure 2B). The top 20 differentially expressed genes between these 3 clusters is depicted in the heatmap in Supplementary Figure 1C, demonstrating that gene expression between these three clusters was highly distinct. A distinguishing feature of the smallest IM cluster was high *Retnla*(Nair et al., 2009a) gene expression, therefore this cluster was termed *Retnla+* macrophages. The remaining macrophage clusters were identified as MHCII hi and MHCII lo IMs. After clustering the cells, hash-tagged cells were sub-setted from the data, keeping the same UMAP-co-ordinates. The purpose of this was to observe where each hash-tagged (i.e. CCR1 and/or CCR5 expressing) population fell within the UMAP and, therefore, what macrophage population they corresponded to (Figure 2Ci). In agreement with previous work(Medina-Ruiz et al., 2024), classical monocytes were almost entirely negative for CCR1 and CCR5 expression. The largest hash-tagged population mapping to the *Retnla+* macrophage cluster was the CCR1+CCR5+ population. Interestingly, CCR1 and CCR5 expression was largely uniform across MHCII hi and MHCII lo macrophages, with the majority of cells in both clusters being CCR1− CCR5+ (Figure 2Ci). Figure 2Cii presents a bar graph depicting the percentage of hash-tagged populations mapping to each cluster. This demonstrates that 86.6% of hash-tagged *Retnla+* macrophages were positive for CCR5. Similar proportions of CCR5+ cells were observed in the hash-tagged MHCII hi macrophage cluster (85.2% CCR5+) and hash-tagged MHCII lo macrophage cluster (86.9% CCR5+). In contrast, 60% of *Retnla+* macrophages were positive for CCR1 whereas only 30% of MHCII hi macrophages and 32% of MHCII lo macrophages were positive for CCR1. Thus, CCR1 and CCR5 expression distributes non-randomly across the 3 identified macrophage clusters.

We next examined the effect of layering *Ccr2* expression onto the hash-tagged populations. Hash-tagging was not required for CCR2 as reporter protein levels correlate well with transcript levels(Medina-Ruiz et al., 2022). We looked at *Ccr2* gene expression across the hash-tagged populations depicted in Figure 2Ci. A cell was considered positive for *Ccr2* if relative expression was >1. As expected, we observed high *Ccr2* expression in the CCR1− CCR5− hash-tagged population, which primarily mapped to classical monocytes (Figure 2Di). A minority (11.7%) of macrophages were also CCR1-CCR5− and, of these, 85% expressed *Ccr2*. CCR2+CCR1-CCR5− cells made up 3.33% of *Retnla+* macrophages, 10.7% of MHCII hi IMs and 10.4% of MHCII lo IMs, respectively. In the CCR1-CCR5+ hash-tagged population, *Ccr2* was expressed in *Retnla+* macrophages and MHCII hi IMs but largely absent from MHCII lo IMs (Figure 2Di). CCR2+CCR1-CCR5+ cells made up 35% of *Retnla+* macrophages, 40.4% of MHCII hi IMs but only 2.7% of MHCII lo macrophages (Figure 2Di). CCR1+CCR5− cells were rare and made up only 2.8% of total hash-tagged cells. CCR2+CCR1+CCR5− cells mapped primarily to *Retnla+* macrophages, making up 10% of the *Retnla+* macrophage population (Figure 2Di). Similar to the CCR1-CCR5+ population, *Ccr2* was expressed in CCR1+CCR5+ cells mapping to *Retnla+* macrophages and MHCII hi IMs but was absent from MHCII lo IMs. *Retnla+* macrophages had the highest proportion of CCR2+CCR1+CCR5+ cells (46.7%), while 21.3% of MHCII hi IMs and 5.8% of MHCII lo IMs were CCR2+CCR1+CCR5+. Looking at the gene expression level, we found that the log2fold change of *Ccr2* was 0.611 in classical monocytes (p value = 1.03×10^−20^) and 0.579 in *Retnla+* macrophages (p value = 4.72×10^−6^), compared to all other clusters. *Ccr2* was not a significant marker gene in MHCII hi or MHCII lo IMs (Supplementary Table 2). The combinatorial expression of *Ccr2*, CCR1 and CCR5 by the three observed macrophage populations is summarized in Figure 2Dii.

Due to *Retnla+* macrophages having the highest proportion of CCR1+CCR2+CCR5+, we hypothesised that they, in fact, correlate to the CCR1+CCR2+CCR5+ MHCII hi macrophage population described in the bulk RNA-sequencing (Figure 1), potentially explaining why the gene expression pattern of CCR1+CCR2+CCR5+ MHCII hi IMs was highly distinct (Figure 1B i&ii). To test this, we used the function FIndAllMarkers in Seurat to identify genes expressing cell surface molecules that could be used to identify *Retnla*+ macrophages by flow cytometry. This was necessary since RELMα, encoded by *Retnla*, is not a cell surface molecule. CD2 and CD226 were selected as alternative markers of *Retnla*+ macrophages for flow cytometry analysis. The specific expression of these genes in the single cell sequencing data is presented in the FeaturePlots shown in Figure 2Ei, demonstrating that expression is highly specific to *Retnla+* macrophages. Lung IMs were stained for flow cytometry analysis (gating strategy provided in Supplementary Figure 2A and Figure 2Eii), which confirmed that CD2+ CD226+ macrophages (i.e. *Retnla+* macrophages) had high MHCII expression and were primarily CCR1+CCR2+CCR5+ (Figure 2Eii). Supplementary Figure 2B shows expression of key *Retnla+* macrophage marker genes identified by single cell sequencing, by the macrophage populations analysed by bulk sequencing (Figure 1). The marker genes were selected first by using the FindAllMarkers function in Seurat. Significant genes correlating to *Retnla+* macrophages were then sorted based on the difference between pct.1 and pct.2. The top 30 of these genes was then sorted by highest to lowest log2fold change. Using this method, the top 14 *Retnla+* macrophage marker genes, identified from the single cell sequencing data set, are depicted in Supplementary Figure 2B, with the addition of *Ccr2*. Importantly, these data demonstrate that the majority of these *Retnla+* macrophage marker genes are preferentially expressed by MHCII hi CCR1+CCR2+CCR5+ve macrophages analysed by bulk RNA-seq, confirming the transcriptional relatedness of these populations.

To summarise, the CCR1+CCR2+CCR5+ MHCII hi IM population described in the bulk RNA-seq analysis (Figure 1) are a distinct population of lung IMs which highly express *Retnla* and likely have a unique functional role in the lung. MHCII hi and MHCII lo IMs differ in their iCCR expression patterns; MHCII hi IMs are predominantly CCR1-CCR2+CCR5+, whereas MHCII lo IMs are predominantly CCR1-CCR2-CCR5+, and express very little *Ccr2* compared to MHCII hi IMs, confirming the bulk RNA-sequencing results. Together, the bulk and single cell RNA-sequencing confirm that iCCR expression is specifically patterned across macrophage subsets in the lung and is not random or chaotic.

### Pseudotime analysis reveals that IMs develop along 2 distinct lineages

In order to understand if changes in IM iCCR expression patterns were altered along a trajectory or if they delineated unrelated macrophage populations, we performed pseudotime analysis using monocle3(Qiu et al., 2017). As it is widely reported that IMs are monocyte derived(Schyns et al., 2018; T’Jonck and Bain, 2023), classical monocytes were selected as the root of the trajectory. These results confirmed that the first macrophage population to arise from monocytes along this trajectory were *Retnla+* macrophages (Figure 3A), the largest fraction of which are CCR2+CCR1+CCR5+. The trajectory then branched and created two major lineages that corresponded to MHCII lo IMs (Figure 3Bi) and MHCII hi IMs (Figure 3Bii). The top 5 differentially expressed genes across pseudotime were calculated for the MHCII lo and MHCII hi trajectories (Supplementary Figure 3A). As cells differentiate into MHCII lo IM they up-regulate genes associated with an M2 macrophage phenotype, including *Cfh*, *Fcna, Marco* and *Mrc1*. In contrast, as cells differentiate into MHCII hi IM they down regulate *Fn1*, *Samhd1* and *Thbs1* and upregulate *Rgs1*, suggesting these cells are digressing from an M2 phenotype to become more inflammatory.

**Figure 3:**
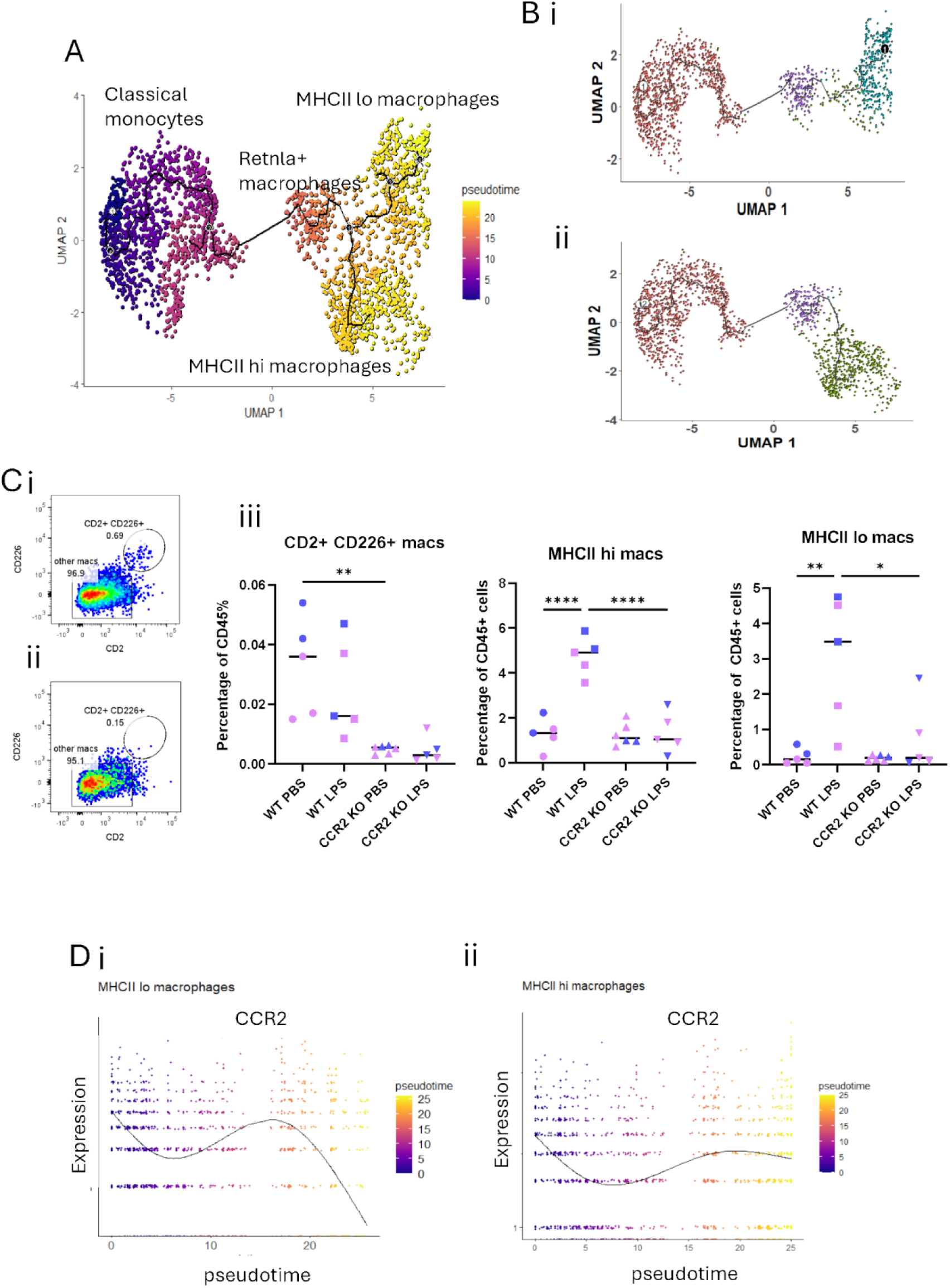
Pseudotime analysis of lung monocyte and macrophage populations demonstrates two distinct macrophages lineages. (A) UMAP plot demonstrating the trajectory from monocytes to macrophages. Colours represent the stage of pseudotime: purple=early in the trajectory, yellow=late in the trajectory. The clusters identified in Figure 2B are labelled. (B) (i) UMAP showing the MHCII lo macrophage lineage trajectory. (ii) UMAP showing the MHCII hi macrophage lineage trajectory. (C) (i) Representative flow cytometry plot showing CD2+ CD226+ expression on F4/80+ macrophages in WT mice. (ii) Representative flow cytometry plot showing CD2+ CD226+ expression of F4/80+ macrophages in CCR2 KO mice. (iii) Quantification of CD2+ CD226+ macrophages (representing *Retnla+* macrophages), MHCII hi macrophages and MHCII lo macrophages in WT and CCR2 KO mice at rest and during LPS induced inflammation. Data are expressed as percentage of CD45+ cells. Colour represents the sex: pink=female mice, blue=male mice. Data were normally distributed and analysed by one-way ANOVA. N= 5-6. *p<0.5, ** p<0.01, ****p<0.0001. (D) (i) *Ccr2* expression across pseudotime in the MHCII lo lineage. (ii) C*cr2* gene expression across pseudotime in the MHCII hi lineage.

Flow cytometry was employed to confirm that *Retnla+* macrophages do indeed derive from monocytes as the pseudotime analysis suggests. To assess this WT mice, and CCR2 KO mice (which display profound monocytopenia in the circulation), were treated intranasally with LPS to induce inflammation in the lung and drive monocyte recruitment. In keeping with their monocytic origin, whilst *Retnla+* macrophages (CD2+ CD226+) were easily detectable in resting and inflamed WT lungs, they were almost entirely absent from resting and inflamed lungs of CCR2 KO mice (Figure 3Ci and ii). Interestingly, and in contrast to MHCII hi and lo macrophages, LPS did not increase *Retnla+* macrophage numbers in WT lungs suggesting that differentiation into *Retnla+* macrophages is an “at rest” process and does not occur in response to inflammatory stimuli (Figure 3Ciii). This is in keeping with the immunoregulatory role of RELMα in the lung that has previously been reported(Lee et al., 2014; Nair et al., 2009b; Sutherland et al., 2018).

Due to the changes observed in *Ccr2* expression between IM subsets, presented in Figure 2D, we measured *Ccr2* across pseudotime for the two observed lineages. As expected, we found that *Ccr2* expression peaked in *Retnla+* macrophages then dramatically decreased across pseudotime along the MHCII lo lineage (Figure 3Di). Although *Ccr2* expression did decrease across pseudotime in the MHCII hi lineage (Figure 3Dii), it was to a much lesser extent than the MHCII lo lineage. It is interesting to note that *Ccr2* expression is downregulated as monocytes differentiate into IMs and is then upregulated again by *Retnla+* macrophages early in the trajectory. This suggests that *Retnla+* macrophages, and MHCII hi macrophages, have a specific biological requirement for CCR2 in addition to CCR1 and CCR5.

Overall, these data demonstrate that the *Retnla+* macrophages expressing all 3 iCCRs sit on a trajectory intermediate between monocytes and terminally differentiated MHCII hi and MHCII lo IMs.

### iCCR expression on macrophages can be patterned by inflammatory mediator exposure

To test the hypothesis that the unique iCCR patterning observed on IM populations is related to discrete functionality of macrophages, indicating non-redundant iCCR expression, we generated BMDM from REP mice and treated them for 24 hours with a range of toll-like receptor ligands representative of a variety of pathogens and inflammatory contexts. Flow cytometric analysis (gating strategy in Supplementary Figure 4) of treated cultures demonstrated that CCR1 expression was increased in response to TLR1/2, TLR4 and TLR5 ligands, but not TLR3 or TLR2/6 ligands, suggestive of regulation by bacterial, but not viral, pathogens (Figure 4Ai). In contrast, CCR2 (Figure 4A ii) and CCR5 (Figure 4Aiii) expression were not significantly increased in response to any TLR ligand. The tailored response of CCR1 to Toll-like receptors signalling in response to bacterial associated ligands, suggests a non-redundant role for CCR1 expression on macrophages.

**Figure 4:**
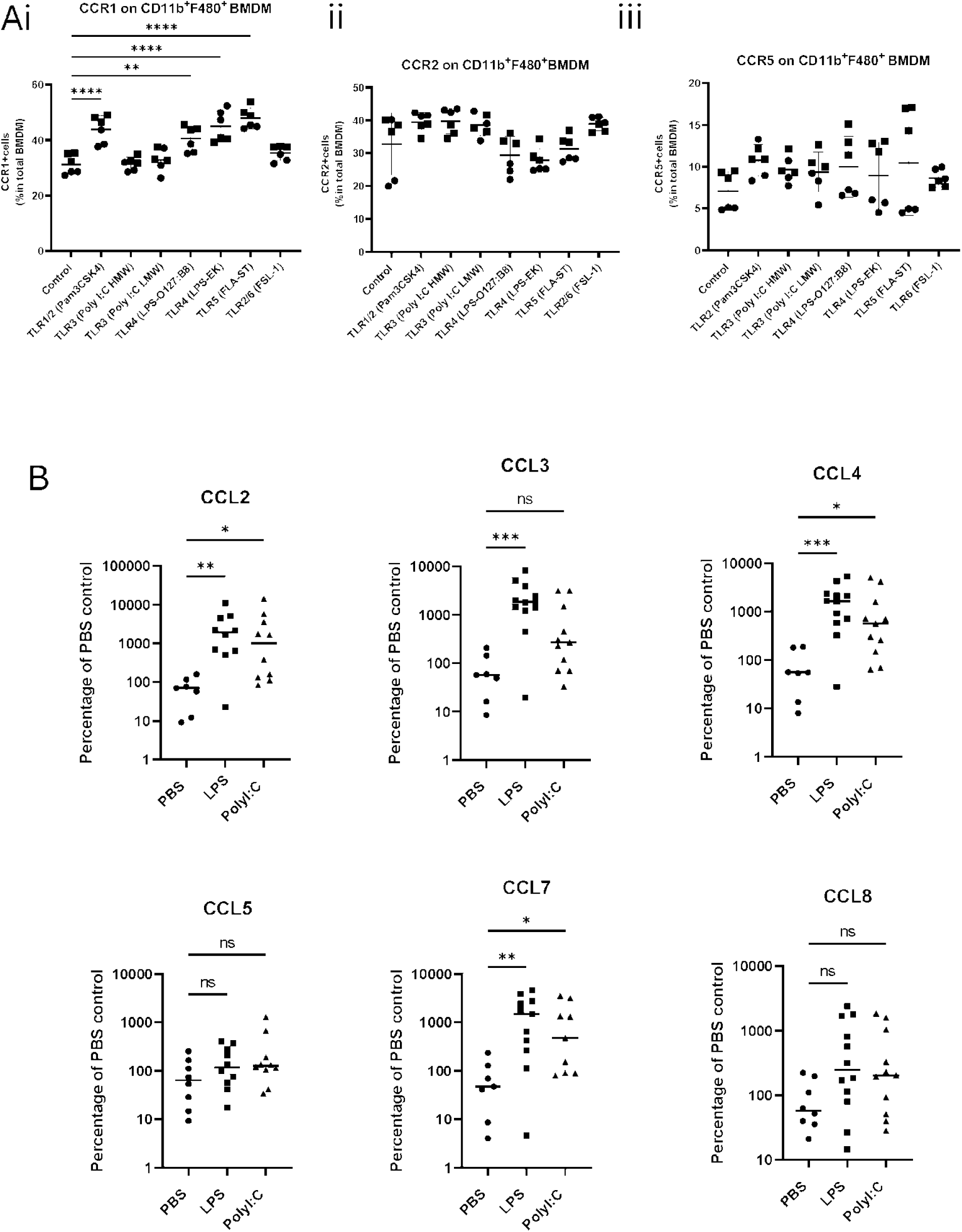
Inflammatory stimuli alter iCCR and ligand expression on macrophages. (A) iCCR expression on BMDM in vitro after 24h inflammatory mediator stimulation. BMDM were first gated based on viability and CD11b and F4/80 expression (gating strategy provided in Supplementary Figure 4A) (i) Percentage of total F4/80+ BMDM expressing CCR1 after 24 hours TLR ligand stimulation. (ii) Percentage of total F4/80+ BMDM expressing CCR2 after 24 hours TLR ligand stimulation. (iii) Percentage of total F4/80+ BMDM expressing CCR5 after 24 hours TLR ligand stimulation. Data in A are pooled from 2 individual experiments denoted by symbol shape. Data were normally distributed and analysed by one-way ANOVA. N=6 *p<0.05, **p<0.01, ***p<0.001, ****p<0.0001. (B) CCL gene expression in the lung 4 hours post intranasal treatment with TLR ligands, LPS and Poly(I:C). CCLs were measured using TaqMan array cards. Delta-delta CT values of target genes were calculated toward 18s RNA and then presented as percentage of PBS control. Data were non-normally distributed and analysed by Kruskal-Wallis test, comparing each treatment condition to the PBS control. N=7-10. *p<0.05, **p<0.01, ***p<0.001.

### Inflammatory chemokine induction in inflamed lung is differentially regulated by bacterial and viral mimetics

To complement the receptor analyses we next examined inflammatory chemokine expression in the lung in response to inhaled TLR ligands indicative of bacterial (LPS) and viral (poly-(I:C)) pathogens. Lungs were harvested 4 hours after TLR ligand treatment and chemokine transcript levels in lung tissue assessed using TaqMan Array cards. These studies focus on CCL2, 3, 4, 5, 7 and 8 expression, as these chemokines are the most highly expressed in the inflamed lung (data not shown). As shown in Figure 4B, with the exception of CCL5 and CCL8, each of these ligands is robustly induced in response to LPS. Notably, whilst CCL2, 4 and 7 are also strongly induced in response to poly(I:C), CCL3 is significantly increased only in response to LPS. As CCL3 is a ligand for CCR1 which, as shown in Figure 4A, is significantly upregulated on BMDM in response to TLR ligands associated with bacterial infection, it is likely that CCL3 signalling via CCR1 has a preferential role in the macrophage response to bacterial infection.

To understand if CCL expression observed in response to LPS and poly(I:C) was tissue specific, or representative of a more general response, we generated data examining chemokine regulation following intradermal injection of the same agents (Figure 5). Expression of CCL2, CCL3, CCL4 and CCL7 were highly similar in the skin compared to the lung in response to both TLR ligands.

**Figure 5:**
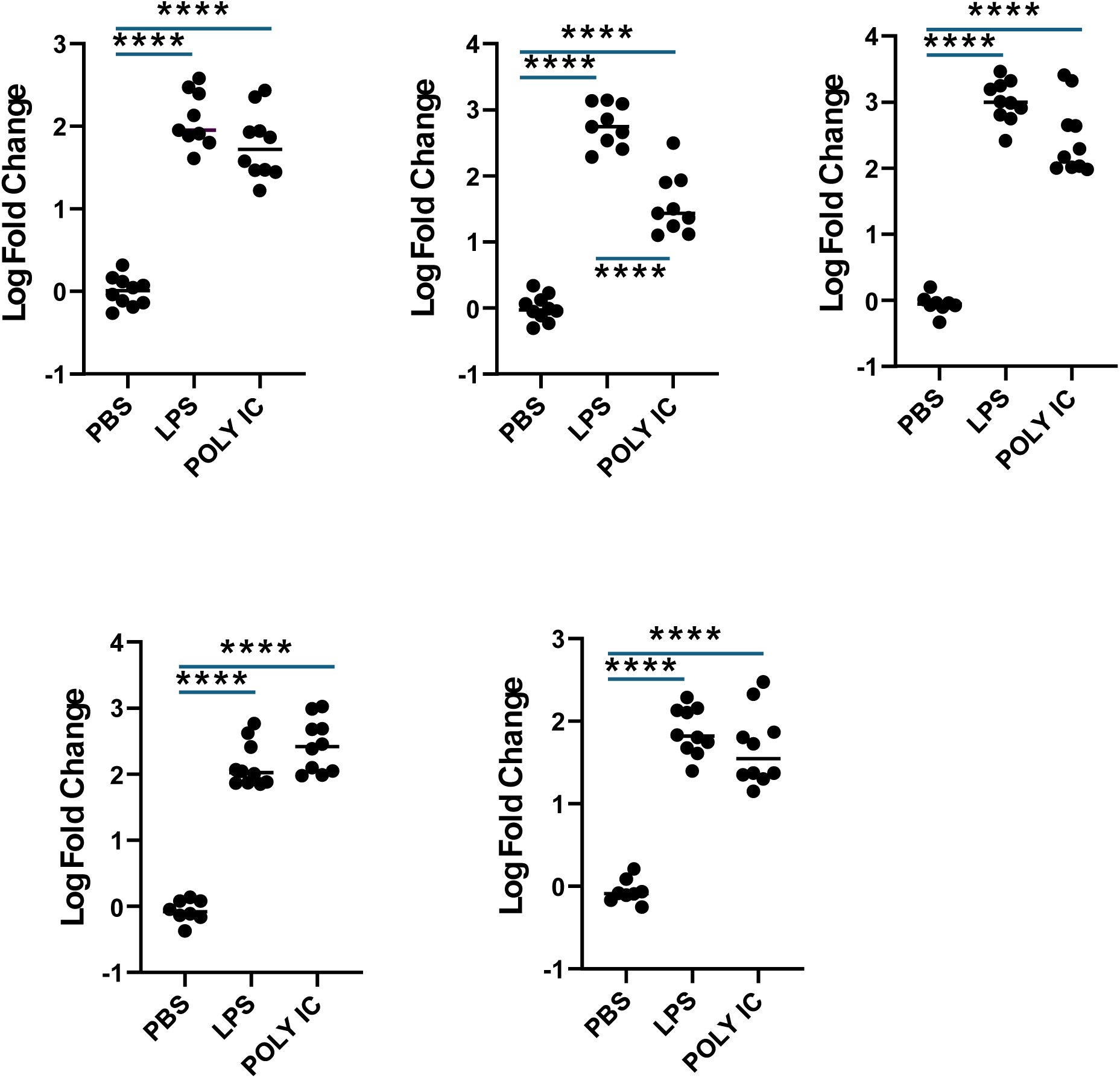
Expression of CCLs in the skin 4 hours after intradermal injection with inflammatory mediators. CCL gene expression in MOUSE dorsal skin 4 hours following intradermal injection with LPS or Poly(I:C). Data were analysed by one-way ANOVA. N= 7-10. ****p<0.0001

To summarise, increased expression of CCL3 is preferentially observed in response to bacterial but not viral agents after both pulmonary and cutaneous administration. The specific increase in CCL3 and CCR1 in response to bacterial ligands, supports the sequencing data presented in Figures 1 and 2, which suggested a non-redundant role for iCCRs in lung IM populations based on non-chaotic expression patterns across IM subsets and significant differential gene expression between IM subsets expressing different iCCR combinations.

## DISCUSSION

For many years it has been argued that, particularly on inflammatory cells, chemokine receptors function in a redundant manner by providing ‘fall-back’ options to protect the robustness of the inflammatory response(Mantovani, 1999; Mantovani, 2018). However, this notion has not been rigorously tested. Here we have examined this focusing on a tight genomic locus incorporating 4 chemokine receptors which we refer to as iCCRs. These receptors have been reported to be expressed in multiple combinations on inflammatory cells and have therefore been suggested to exemplify the notion of redundancy. Using multi-chemokine receptor reporter mice(Medina-Ruiz et al., 2022), we have previously confirmed that macrophages can indeed display essentially all combinations of CCR1, CCR2 and CCR5 but determining the relevance of this for redundancy has been difficult. In part this is because clear roles for CCR1 and CCR5, in macrophage biology, have not yet been defined(Dyer et al., 2019) and thus determining potential functional redundancy is problematic. Here we have used transcriptomic approaches to test the hypothesis that chemokine receptors do not function in a redundant manner and that macrophages expressing different iCCR combinations represent distinct macrophage populations.

Using both bulk and single cell RNA sequencing we have demonstrated that, for both MHCII hi and MHCII lo IMs, cells expressing different iCCR combinations are transcriptionally distinct. Furthermore, bioinformatic analysis of single cell RNA-Seq data shows that chemokine receptor expression can be used to define a trajectory from CCR2+ve inflammatory monocytes, via a *Retnla*+ intermediate macrophage population co-expressing CCR1, CCR2 and CCR5, to mature macrophages expressing various combinations of CCR1, CCR2 and CCR5. Notably, CCR5 dominates receptor expression patterns in MHCII hi and lo macrophages frequently in combination with CCR2 in MHCII hi cells. Some expression of CCR1 is noted but it is limited. Overall, these data suggest that macrophage differentiation from inflammatory monocytes proceeds via a *Retnla*+ intermediate population expressing all 3 iCCRs and that expression is subsequently refined, and restricted, on fully mature macrophages. These data are from IMs derived from resting lungs and suggest that the predominant default in mature cells is CCR5 expression. Expression of multiple cell surface proteins in precursor cells, with progressive restriction upon differentiation, is also seen in other situations(Tan et al., 2015; Wada et al., 2018) suggesting that this may be a more general phenomenon in biology. Furthermore, the chemokine receptor CX3CR1 has been reported to mark selective differentiation states of human and murine T cells(Zwijnenburg et al., 2023) suggesting that chemokine receptors, as markers of lineage differentiation, may have broader applicability.

We propose that the Retnla+ population represents a pre-mature macrophage population and that multiple iCCR expression equips these cells with the ability to rapidly migrate towards a range of possible pathogens, or tissue damage, potentially associated with alternative chemokine ligand expression. We further hypothesise that iCCR expression is then progressively restricted in the macrophages in response to sensing of the particular inflammatory environment, thus aligning iCCR expression with pathogen challenge. In addition, our data show that bacterial pathogen mimetics i.e. TLR ligands can skew macrophage iCCR expression in favour of CCR1 suggesting that final iCCR expression patterns can be manipulated by the inflammatory environment. In terms of CCR1, it is of note that CCL3 is more strongly induced by LPS than poly(I:C) suggesting that this chemokine may be preferentially important for antibacterial responses. Importantly, CCL3 has been shown, in a range of inflammatory situations, to be a physiological substrate for the enzyme DPPIV (CD26)(Proost et al., 2000). This enzyme (expressed in the lung: https://www.gtexportal.org/home/gene/DPP4 and data not shown) cleaves the 2 most N-terminal amino acids from CCL3, changing its receptor-binding patterns from similar affinities for CCR1 and CCR5 to predominantly CCR1 binding(Mortier et al., 2016; Nibbs et al., 1999). Thus, the ability of LPS to induce both CCR1 in macrophages, and CCL3 at inflamed sites, aligns macrophage chemotactic responsiveness to pathogen-specific chemokine expression. Further investigation is required to try to align alternative pathogen challenges with specific iCCR expression changes and associated ligand profiles.

In summary therefore we have used transcriptomic approaches to demonstrate that inflammatory chemokine receptor expression in macrophages progresses from multi-receptor expressing macrophages to more restricted expression in mature cells. Our data therefore confirm multiple iCCR expression on macrophages and demonstrate that this is not a consequence of redundancy but that individual receptor expression patterns delineate transcriptionally distinct cells on the monocyte to mature-macrophage trajectory.

## MATERIALS AND METHODS

### Animals

CCR2 KO (Jackson labs) (C57BL/6J background) and REP(Medina-Ruiz et al., 2022) (C57BL/6N background) mice were housed in a specific pathogen-free animal facility at the University of Glasgow. WT controls were matched to the appropriate background of the transgenic mice. All mice were aged between 7-12 weeks, using a mixture of male and female mice for most experiments. Only female mice were used for single cell RNA-seq as samples were combined into a single pool. Procedures were approved by the local University of Glasgow ethics committee and animal experimentation was carried out under a UK Home Office Licence in accordance with the revised Animal (Scientific Procedures) Act 1986.

### Bulk RNA-Sequencing

Lungs were digested using Liberase and processed into a single cell suspension as described above. Cells were incubated for 20 minutes at 4⁰C with efluor 506 viability stain (Thermo Fisher Scientific, 65-0866-14), diluted 1:1000 in PBS. Fc receptor binding was blocked by incubating the cells in FcR blocking reagent (Miltenyi Biotec, 130-092-575) Staining was performed using the following antibodies: anti-mouse CD45.2 PerCP-Cy5.5 (BioLegend; clone 104; cat. no 109828), anti-mouse/human CD11b APC-Cy7 (Biolegend; clone M1/70; cat no. 101226), anti-mouse CD64 Brilliant Violet 786 (BD Biosciences; clone: X54-5/7.1; cat.no. 569507), anti-mouse MHCII Brilliant Violet 605 (BioLegend; clone: M5/114.15.2, cat.no. 107639), anti-mouse F4/80 PE-Cy7 (Thermo Fisher; clone: BM8; cat no. 25-4801-82), anti-mouse Ly6C Alexa Fluor 700 (BioLegend; clone: HK1.4; cat.no. 128024), anti-mouse Ly6G Alexa Fluor 700 (BioLegend; clone: 1A8; cat no. 127622) and anti-mouse SiglecF Alexa Fluor 700 (BioLegend; clone: S17007L; cat.no. 155534). MHCII hi and MHCII lo macrophages expressing various combinations of iCCRs were sorted on the BD FACS Aria using the gating strategies presented in Supplementary Figure 1A and Figure 1A. Cells were collected into RLT buffer (Qiagen, 79216) containing 10μl/ml of 2-ME and stored at −80°C for RNA extraction.

RNA extraction, library preparation and bulk RNA-sequencing was carried out as previously described(Medina-Ruiz et al., 2024). Briefly, RNA was extracted using an RNeasy Micro Kit (Qiagen) as per the manufacturer’s instructions and mRNA libraries were prepared using the NEBNext Single Cell/Low Input RNA Library Prep Kit for Illumina (New England Biolabs; cat. no. E6420L). Samples were sequenced using paired-end sequencing on the NextSeq2000 sequencing platform (Illumina) (40 million reads sequencing depth). FastQ files were assessed using FastP(Chen et al., 2018), and aligned to the mouse reference genome (GRCm38.91) using STAR (2.7.10a)(Dobin et al., 2013). The expression and differential expression values were generated using DESeq2 (version 1.24)(Love et al., 2014) or differential comparisons, an A versus B model with no additional covariates was used. All other parameters were left to default. The processed data were then visualized using Searchlight(Cole et al., 2021), specifying one differential expression workflow for each comparison, an absolute log2 fold cutoff of 1, and adjusted p value of 0.05.

### Sample Processing for Single-Cell RNA-Sequencing

Lungs from 3x female REP mice were perfused with 20 ml PBS and digested in 0.44 Wunsch units of Liberase (Merck, LIBTM-RO) at 37⁰C for 1 hour. Digested samples were passed through a 70µm filter to obtain a single cell suspension and pooled together for sorting. Cells were stained first with efluor 506 viability stain (Thermo Fisher Scientific, 65-0866-14), then with the following antibodies: anti-mouse/human CD11b PE-Cy7 (BioLegend; clone M1/70; cat no. 101216), anti-mouse Ly6G Brilliant Violet 711 (BioLegend; clone 1A8; cat no. 127643) and anti-mouse SiglecF Brilliant Violet 711 (BD Biosciences; clone E50-2440; cat no. 740764). CD11b+ Ly6G− SiglecF− cells were sorted from total viable cells using the BD FACSAria and collected into 1% BSA in PBS. Four more samples were sorted from the CD11b+ Ly6G− SigF-gate based on iCCR expression (CCR1-CCR5+, CCR1+CCR5+, CCR1+CCR5-, CCR1-CCR5-); samples sorted based on iCCR expression were hash-tagged using Total-SeqA antibodies (BioLegend; all antibodies were a combination of clones M1/42 and 30-F11 conjugated to different hashtag oligo sequences, cat numbers = 155801, 155803, 155805, 155811) and pooled prior to library generation. A separate library was generated for the CD11b+ Ly6G− SiglecF− sample.

### Single Cell RNA-Sequencing and Analysis

Single-cell libraries were generated and sequenced as previously described(Medina-Ruiz et al., 2024). Reads were aligned using the count function in 10X Genomics Cell Ranger, and outputs were imported into R (version 4.3.1) for downstream analysis using Seurat, version 4.4.0(Butler et al., 2018; Hao et al., 2021; Hao et al., 2024; Satija et al., 2015; Stuart et al., 2019). Single cell hash-tagged data was demultiplexed using the package CellhashR(Boggy et al., 2022). The data were quality controlled by adjusting the range of features and counts included in downstream analysis. the range of features included in downstream analysis was 100–4000, and the range of counts was 100–25,000. In addition, cells were removed if they did not meet the threshold for mitochondrial percentage (<5%). Data were normalized using both log normalization and SC-Transform. Integration features were identified using default features of the Seurat function SelectIntegrationFeatures. The function PrepSCTIntegration was then run on the SCT assay using the selected integration features. Thirty principle-components (PCs) were included in the function FindIntegrationAnchors to identify the anchors required for integration. Thirty PCs were selected based on the results of an Elbow Plot displaying PCs 1-50 (data not shown). Finally, data were integrated using the Seurat function IntegrateData, again including 30 PCs. Seurat was updated to version 5 to run downstream analysis. Data was plotted and clustered using the functions FindVariableFeatures, ScaleData, RunPCA (npcs = 10), RunUMAP (reduction = “pca”, dims = 1:10), FindNeighbours (reduction = “pca”, dims = 1:10 and FindClusters (resolution = 0.5). The effect of cell-cycle was assessed using the function CellCycleScoring and the cell cycle score was regressed from the data for downstream analysis. Uniform Manifold Approximation and Projection (UMAP) dimensionality reduction was applied for visualization of the data in Dimensional Reduction Plots and Feature Plots. Pseudotime analysis was performed in R (version 4.3.1) using Monocle 3(Qiu et al., 2017; Trapnell et al., 2014). Briefly, the integrated Seurat object was transformed into a Cell Data Set (CDS) object using the default parameters of the function as.cell_data_set. Pre-processing of the CDS object was then carried out using the function preprocess_cds (num_dim = 100). Seurat clusters from previous analysis were applied to the CDS object. Pseudotime analysis was then performed using the functions: learn_graph (use_partitions = TRUE), order_cells (reduction method = UMAP) and estimate_size_factors.

### Flow cytometry

#### Lung Macrophages

Mice were culled via CO_2_ inhalation and lungs perfused with 20ml PBS. Lungs were processed into a single cell suspension as described above. Samples were then stained with efluor 506 viability stain, diluted 1:1000 in PBS. Fc receptor binding was blocked by incubating the cells in FcR blocking reagent (Miltenyi Biotec; cat.no. 130-092-575). For experiments with CCR2 KO mice and equivalent WT controls, staining was performed using the following antibodies: anti-mouse CD45.2 – APC-Fire 750 (BioLegend; clone: 104; cat.no. 109852), anti-mouse Ly6G – Brilliant Violet 711 (BioLegend; clone: 1A8; cat.no. 127643), anti-mouse Ly6C – Alexa Fluor 488 (BioLegend, clone: HK1.4; cat.no. 128022), SiglecF – Brilliant Ultra-violet 737 (BD Biosciences; clone: E50-2440; 570753), F4/80 – PE (BioLegend; clone: W20065B; cat. no. 111604), anti-mouse CD11b – PE-Cy7 (BioLegend; clone M1/70), anti-mouse CD64 – Brilliant Violet 786 (BD Biosciences; clone: X54-5/7.1; cat.no. 569507)), anti-mouse MHCII Brilliant Violet 605 (BioLegend; clone: M5/114.15.2, cat.no. 107639), CD11c – Alexa Fluor 700 (BD Biosciences; clone: HL3; cat.no. 560583), anti-mouse CD2 – APC (BioLegend; clone: RM2-5; cat. no. 100112), and anti-mouse CD226 – PerCP-Cy5.5 (BioLegend; clone: 10E5; cat. no. 128814). For experiments using REP mice, antibodies targeting Ly6G, MHCII and CD226 were the same as stated above for the CCR2 KO experiment. Additional antibodies used were: anti-mouse CD45 APC-Cy7 (BioLegend, clone: 30-F11; cat. no. 103116), anti-mouse/human CD11b Brilliant Ultra Violet 805 (BD Biosciences; clone: M1/70; cat.no. 568345), anti-mouse SiglecF Brilliant Violet 711 (BD Biosciences; clone: E50-2440; cat no. 740764), anti-mouse F4/80 Brilliant Violet 650 (BioLegend; clone: BM8; cat. no. 123149) and anti-mouse CD2 PE-Cy7 (BioLegend; clone RM2-5; cat.no. 100113). Antibody staining was performed at 4⁰C for 20 minutes. Samples were acquired using BD Fortessa and analysed using FlowJo (v10).

#### Bone marrow derived macrophages

BMDM were cultured and treated as described below, then lifted from plastic using Trypsin-EDTA (0.5%) (Thermo Fisher Scientific, 25300054) and stained with viability stain eFluor 506 diluted 1:1000 in PBS. Fc receptor binding was blocked by incubating the cells in FcR blocking reagent and staining was performed by incubating cells with the following cocktail of antibodies for 20 minutes at 4⁰C: CD11b – APC-Cy7 (Biolegend, clone = M1/70) and F4/80 – Brilliant Violet 785 (Biolegend, clone = BM8). Samples were acquired using BD Fortessa and analysed using FlowJo (v10).

### Culture of bone marrow derived macrophages and treatment with inflammatory mediators

Bone marrow derived macrophages were cultured as previously described(Bartolini et al., 2022). Briefly, Tibia and femurs were collected from culled mice and bone marrow (BM) was flushed out, strained through a 70μm cell strainer and red cells lysed using ACK lysis buffer (Gibco, A1049201), to obtain a single-cell suspension of leukocytes. Cells suspended in Glasgow’s Minimal Essential Medium (Gibco, 11710035 supplemented with 15% L929 conditioned media, 10% FBS (Thermo Fisher, 10500064), 1% l-glutamine (Thermo Fisher, 25030024), 100mM Sodium Pyruvate solution (Sigma, S8636), 1% MEM nonessential amino acids (Gibco, 1140-035), 50 μM b-ME (Gibco, 31350-010), and 0.2% primocin (InvivoGen, ant-pm-1) were seeded at 1.2 ×10^6^ cells per well of a 6-well plate. Cells were cultured for 7 days, to allow macrophages to differentiate, and treated with Toll-like Receptor (TLR) agonists for the final 24 hours. TLR agonists used were: Pam3CSK4 (TLR1/2 agonist used at 150ng/ml; InvivoGen; tlrl-pms), Poly(I:C) (HMW) (TLR3 agonist used at 1µg/ml; InvivoGen; tlrl-hklm), Poly(I:C) (LMW) (TLR3 agonist used at 5µg/ml; InvivoGen; tlrl-picw), LPS-O127:B8 (TLR4 agonist used at 100ng/ml; Sigma Aldrich; L45-16), LPS-EK (TLR4 agonist used at 5µg/ml; InvivoGen; tlrl-eklps), FLA-ST (TLR5 agonist used at 5µg/ml; InvivoGen; tlrl-stfla), FSL-1 (TLR2/6 agonist used at 50ng/ml; InvivoGen; tlrl-fsl).

### Intranasal inflammatory mediator treatments

Mice were anaesthetised using inhaled isofluorane (4% v/v + 2L O2/min). and treated intranasally with 25µl LPS (1000ug/ml) (Invitrogen, 00-4976-03, 25µl PolyI:C (1000ug/ml) (Invivogen, tlrl-pic), or 25µl PBS as a control. Four hours later, mice were culled via lethal intraperitoneal injection of the pentobarbital sodium solution, Dolethal. lungs were perfused with 20ml PBS and collected into in RNAlater Stabilization Solution (Thermo Fisher Scientific, AM7020) for downstream processing.

### Gene expression analysis using TaqMan Array Cards

Tissue samples intended for RNA extraction were stored in RNAlater for at least 24h. Tissues were lysed and whole RNA was extracted using TRIzol™ Reagent (Thermo Fisher Scientific, cat.no.15596026) -chloroform extraction, followed column clean up using PureLink™ RNA Mini Kit (Invitrogen, cat.no. 4387406) with an on-column DNase (Qiagen, 79254) digest. RNA was converted to cDNA using the High-Capacity RNA-to-cDNA™ Kit (Applied Biosystems, cat.no. 10704217). The expression of 48 genes, including housekeeping genes were determined using custom made TaqMan Array Cards (Thermo Fisher Scientific, cat. no. 4342253, design ID RTRWE3D). 1 µg of cDNA with TaqMan™ Fast Advanced Master Mix (ThermoFisher Scientific, cat. no. 4444557) was loaded per slot and array cards were run on a QuantStudio™ 7 Flex (Thermo Fisher Scientific, cat. no. 4485701) real-time PCR system. Delta-delta CT values of target genes were calculated toward 18s RNA.

### Data availability

Raw and processed sequencing data have been uploaded to Gene Expression Omnibus. Single cell RNA-Seq data has the accession number: GSE305101 and bulk RNA-Seq data has the accession number: GSE306514. Original code used to analyse single cell RNA-Seq data was uploaded to Zenodo and can be found at the following: https://doi.org/10.5281/zenodo.16876331

This work was funded by grants from the Wellcome Trust and the Medical Research Council. We gratefully acknowledge the assistance of University of Glasgow MVLS SRF.

## REFERENCES

Bachelerie, F., A. Ben-Baruch, A.M. Burkhardt, C. Combadiere, J.M. Farber, G.J. Graham, R. Horuk, A.H. Sparre-Ulrich, M. Locati, A.D. Luster, A. Mantovani, K. Matsushima, P.M. Murphy, R. Nibbs, H. Nomiyama, C.A. Power, A.E.I. Proudfoot, M.M. Rosenkilde, A. Rot, S. Sozzani, M. Thelen, O. Yoshie, and A. Zlotnik. 2014. International Union of Pharmacology. LXXXIX. Update on the Extended Family of Chemokine Receptors and Introducing a New Nomenclature for Atypical Chemokine Receptors. Pharmacological Reviews 66:1–79.

Bartolini, R., L. Medina-Ruiz, A.J. Hayes, C.J. Kelly, H.A. Halawa, and G.J. Graham. 2022. Inflammatory Chemokine Receptors Support Inflammatory Macrophage and Dendritic Cell Maturation. Immunohorizons 6:743–759.

Boggy, G.J., G.W. McElfresh, E. Mahyari, A.B. Ventura, S.G. Hansen, L.J. Picker, and B.N. Bimber. 2022. BFF and cellhashR: analysis tools for accurate demultiplexing of cell hashing data. Bioinformatics 38:2791–2801.

Butler, A., P. Hoffman, P. Smibert, E. Papalexi, and R. Satija. 2018. Integrating single-cell transcriptomic data across different conditions, technologies, and species. Nature Biotechnology 36:411–420.

Chen, S., Y. Zhou, Y. Chen, and J. Gu. 2018. fastp: an ultra-fast all-in-one FASTQ preprocessor. Bioinformatics 34:i884–i890.

Cole, J.J., B.A. Faydaci, D. McGuinness, R. Shaw, R.A. Maciewicz, N.A. Robertson, and C.S. Goodyear. 2021. Searchlight: automated bulk RNA-seq exploration and visualisation using dynamically generated R scripts. BMC Bioinformatics 22:411.

Dobin, A., C.A. Davis, F. Schlesinger, J. Drenkow, C. Zaleski, S. Jha, P. Batut, M. Chaisson, and T.R. Gingeras. 2013. STAR: ultrafast universal RNA-seq aligner. Bioinformatics 29:15–21.

Dyer, D.P., L. Medina-Ruiz, R. Bartolini, F. Schuette, C.E. Hughes, K. Pallas, F. Vidler, M.K.L. Macleod, C.J. Kelly, K.M. Lee, C.A.H. Hansell, and G.J. Graham. 2019. Chemokine Receptor Redundancy and Specificity Are Context Dependent. Immunity 50:378–389.e375.

Gibbings, S.L., S.M. Thomas, S.M. Atif, A.L. McCubbrey, A.N. Desch, T. Danhorn, S.M. Leach, D.L. Bratton, P.M. Henson, W.J. Janssen, and C.V. Jakubzick. 2017. Three Unique Interstitial Macrophages in the Murine Lung at Steady State. Am J Respir Cell Mol Biol 57:66–76.

Griffith, J.W., C.L. Sokol, and A.D. Luster. 2014. Chemokines and chemokine receptors: positioning cells for host defense and immunity. Annu Rev Immunol 32:659–702.

Hao, Y., S. Hao, E. Andersen-Nissen, W.M. Mauck, S. Zheng, A. Butler, M.J. Lee, A.J. Wilk, C. Darby, M. Zager, P. Hoffman, M. Stoeckius, E. Papalexi, E.P. Mimitou, J. Jain, A. Srivastava, T. Stuart, L.M. Fleming, B. Yeung, A.J. Rogers, J.M. McElrath, C.A. Blish, R. Gottardo, P. Smibert, and R. Satija. 2021. Integrated analysis of multimodal single-cell data. Cell 184:3573–3587.e3529.

Hao, Y., T. Stuart, M.H. Kowalski, S. Choudhary, P. Hoffman, A. Hartman, A. Srivastava, G. Molla, S. Madad, C. Fernandez-Granda, and R. Satija. 2024. Dictionary learning for integrative, multimodal and scalable single-cell analysis. Nature Biotechnology 42:293–304.

Hariharan, D., S.D. Douglas, B. Lee, J.P. Lai, D.E. Campbell, and W.Z. Ho. 1999. Interferon-gamma upregulates CCR5 expression in cord and adult blood mononuclear phagocytes. Blood 93:1137–1144.

Kaufmann, A., R. Salentin, D. Gemsa, and H. Sprenger. 2001. Increase of CCR1 and CCR5 expression and enhanced functional response to MIP-1α during differentiation of human monocytes to macrophages. Journal of Leukocyte Biology 69:248–252.

Lee, M.-R., D. Shim, J. Yoon, H.S. Jang, S.-W. Oh, S.H. Suh, J.-H. Choi, and G.T. Oh. 2014. Retnla Overexpression Attenuates Allergic Inflammation of the Airway. PloS one 9:e112666.

Love, M.I., W. Huber, and S. Anders. 2014. Moderated estimation of fold change and dispersion for RNA-seq data with DESeq2. Genome biology 15:550.

Mantovani, A. 1999. The chemokine system: redundancy for robust outputs. Immunol Today 20:254–257.

Mantovani, A. 2018. Redundancy and robustness versus division of labour and specialization in innate immunity. Semin Immunol 36:28–30.

Medina-Ruiz, L., R. Bartolini, H. Mathie, H.A. Halawa, M. Cunningham, and G.J. Graham. 2024. CCR1 and CCR2 Coexpression on Monocytes Is Nonredundant and Delineates a Distinct Monocyte Subpopulation. The Journal of Immunology ji2400007.

Medina-Ruiz, L., R. Bartolini, G.J. Wilson, D.P. Dyer, F. Vidler, C.E. Hughes, F. Schuette, S. Love, M. Pingen, A.J. Hayes, J. Fu, A.F. Stewart, and G.J. Graham. 2022. Analysis of combinatorial chemokine receptor expression dynamics using multi-receptor reporter mice. eLife 11:e72418.

Mortier, A., M. Gouwy, J. Van Damme, P. Proost, and S. Struyf. 2016. CD26/dipeptidylpeptidase IV-chemokine interactions: double-edged regulation of inflammation and tumor biology. J Leukoc Biol 99:955–969.

Nair, M.G., Y. Du, J.G. Perrigoue, C. Zaph, J.J. Taylor, M. Goldschmidt, G.P. Swain, G.D. Yancopoulos, D.M. Valenzuela, A. Murphy, M. Karow, S. Stevens, E.J. Pearce, and D. Artis. 2009a. Alternatively activated macrophage-derived RELM-{alpha} is a negative regulator of type 2 inflammation in the lung. J Exp Med 206:937–952.

Nair, M.G., Y. Du, J.G. Perrigoue, C. Zaph, J.J. Taylor, M. Goldschmidt, G.P. Swain, G.D. Yancopoulos, D.M. Valenzuela, A. Murphy, M. Karow, S. Stevens, E.J. Pearce, and D. Artis. 2009b. Alternatively activated macrophage-derived RELM-α is a negative regulator of type 2 inflammation in the lung. Journal of Experimental Medicine 206:937–952.

Nibbs, R.J., J. Yang, N.R. Landau, J.H. Mao, and G.J. Graham. 1999. LD78beta, a non-allelic variant of human MIP-1alpha (LD78alpha), has enhanced receptor interactions and potent HIV suppressive activity. J Biol Chem 274:17478–17483.

Proost, P., P. Menten, S. Struyf, E. Schutyser, I. De Meester, and J. Van Damme. 2000. Cleavage by CD26/dipeptidyl peptidase IV converts the chemokine LD78beta into a most efficient monocyte attractant and CCR1 agonist. Blood 96:1674–1680.

Proudfoot, A.E.I. 2002. Chemokine receptors: multifaceted therapeutic targets. Nature Reviews Immunology 2:106–115.

Qiu, X., Q. Mao, Y. Tang, L. Wang, R. Chawla, H.A. Pliner, and C. Trapnell. 2017. Reversed graph embedding resolves complex single-cell trajectories. Nat. Methods 14:979–982.

Rot, A., and U.H. von Andrian. 2004. Chemokines in innate and adaptive host defense: basic chemokinese grammar for immune cells. Annu Rev Immunol 22:891–928.

Satija, R., J.A. Farrell, D. Gennert, A.F. Schier, and A. Regev. 2015. Spatial reconstruction of single-cell gene expression data. Nature Biotechnology 33:495–502.

Schyns, J., F. Bureau, and T. Marichal. 2018. Lung Interstitial Macrophages: Past, Present, and Future. J Immunol Res 2018:5160794.

Stuart, T., A. Butler, P. Hoffman, C. Hafemeister, E. Papalexi, W.M. Mauck, Y. Hao, M. Stoeckius, P. Smibert, and R. Satija. 2019. Comprehensive Integration of Single-Cell Data. Cell 177:1888–1902.e1821.

Sutherland, T.E., D. Rückerl, N. Logan, S. Duncan, T.A. Wynn, and J.E. Allen. 2018. Ym1 induces RELMα and rescues IL-4Rα deficiency in lung repair during nematode infection. PLoS pathogens 14:e1007423.

T’Jonck, W., and C.C. Bain. 2023. The role of monocyte-derived macrophages in the lung: It’s all about context. The International Journal of Biochemistry & Cell Biology 159:106421.

Tan, L., Q. Li, and X.S. Xie. 2015. Olfactory sensory neurons transiently express multiple olfactory receptors during development. Mol Syst Biol 11:844.

Trapnell, C., D. Cacchiarelli, J. Grimsby, P. Pokharel, S. Li, M. Morse, N.J. Lennon, K.J. Livak, T.S. Mikkelsen, and J.L. Rinn. 2014. The dynamics and regulators of cell fate decisions are revealed by pseudotemporal ordering of single cells. Nat Biotechnol 32:381–386.

Wada, T., S. Wallerich, and A. Becskei. 2018. Stochastic Gene Choice during Cellular Differentiation. Cell reports 24:3503–3512.

Wynn, T.A., A. Chawla, and J.W. Pollard. 2013. Macrophage biology in development, homeostasis and disease. Nature 496:445–455.

Zlotnik, A., and O. Yoshie. 2000. Chemokines: a new classification system and their role in immunity. Immunity 12:121–127.

Zwijnenburg, A.J., J. Pokharel, R. Varnaitė, W. Zheng, E. Hoffer, I. Shryki, N.R. Comet, M. Ehrström, S. Gredmark-Russ, L. Eidsmo, and C. Gerlach. 2023. Graded expression of the chemokine receptor CX3CR1 marks differentiation states of human and murine T cells and enables cross-species interpretation. Immunity 56:1955–1974.e1910.

